# Attribution of genetic engineering: A practical and accurate machine-learning toolkit for biosecurity

**DOI:** 10.1101/2020.08.22.262576

**Authors:** Ethan C. Alley, Miles Turpin, Andrew Bo Liu, Taylor Kulp-McDowall, Jacob Swett, Rey Edison, Stephen E. Von Stetina, George M. Church, Kevin M. Esvelt

## Abstract

The promise of biotechnology is tempered by its potential for accidental or deliberate misuse. Reliably identifying telltale signatures characteristic to different genetic designers, termed *genetic engineering attribution*, would deter misuse, yet is still considered unsolved. Here, we show that recurrent neural networks trained on DNA motifs and basic phenotype can reach 70% attribution accuracy distinguishing between over 1,300 labs. To make these models usable in practice, we introduce a framework for weighing predictions against other investigative evidence using calibration, and bring our model to within 1.6% of perfect calibration. Additionally, we demonstrate that simple models can accurately predict both the nation-state-of-origin and ancestor labs, forming the foundation of an integrated attribution toolkit which should promote responsible innovation and international security alike.

## Introduction

After a nearly decade-long, $100 million dollar investigation into the 2001 Amerithrax anthrax attacks^1^, the National Academy of Sciences reported that “it is not possible to reach a definitive conclusion about the origins…based solely on the available scientific evidence”^2^. In the aftermath of Amerithrax, the scientific community mobilized to develop genetic^3^ and phenotypic^4^ methods for forensic attribution of biological attacks. However, these efforts were severely constrained by the limits of biotechnology; satisfactory sequencing of the anthrax agent’s genome would have cost $500,000 dollars at the time^5^. Today, exponential improvement in tools for biotechnology have made this and other life-saving tasks increasingly inexpensive and accessible, yet these advancements have not been matched with corresponding improvements in tools to support responsible innovation in genetic engineering. In particular, attribution is still considered technically challenging and unsolved^6–9^.

Just as programmers leave clues in code, biologists “programming” an engineered organism differ on unique design decisions (e.g. promoter choice), style (e.g. preferred codon optimization), intent (e.g. functional gene selection), and tools (e.g. cloning method). These design elements, together with evolutionary markers, form designer “signatures”. Advances in high-throughput, spatial^12^, and distributed^13^ sequencing^12–14^, and omic-scale phenotyping^15^ make these signatures easier to collect, but require complex data analysis. Recent work suggests that deep learning, a flexible paradigm for statistical models built from differentiable primitives, can facilitate complex biological data analysis^16–21^. We propose deploying these methods to develop a toolkit of machine learning algorithms which can infer the attributes of genetically engineered organisms— like lab-of-origin— to support biotechnology stakeholders and enable ongoing efforts to scale and automate biosecurity^22^.

A prior attempt to use deep learning for genetic engineering attribution provided evidence that it might be possible, but was less than 50% accurate^23^. Here, we reach over 70% lab-of-origin attribution accuracy using a biologically motivated approach based on learned DNA motifs, simple phenotype information, and Recurrent Neural Networks (RNNs) on a model attribution scenario with data from the world’s largest plasmid repository, Addgene^24^ (Fig. 1a). We show that this algorithm, which we call *deteRNNt*, can provide more calibrated probabilities— a prerequisite for practical use— and with simple models demonstrate that a wider range of attribution tools are possible. By simultaneously addressing the need for accuracy, more calibrated uncertainty, and broad capabilities, this computational forensic framework can help enable the characterization of engineered biological materials, promote technology development which is accountable to community stakeholders, and deter misuse.

**Figure 1.**
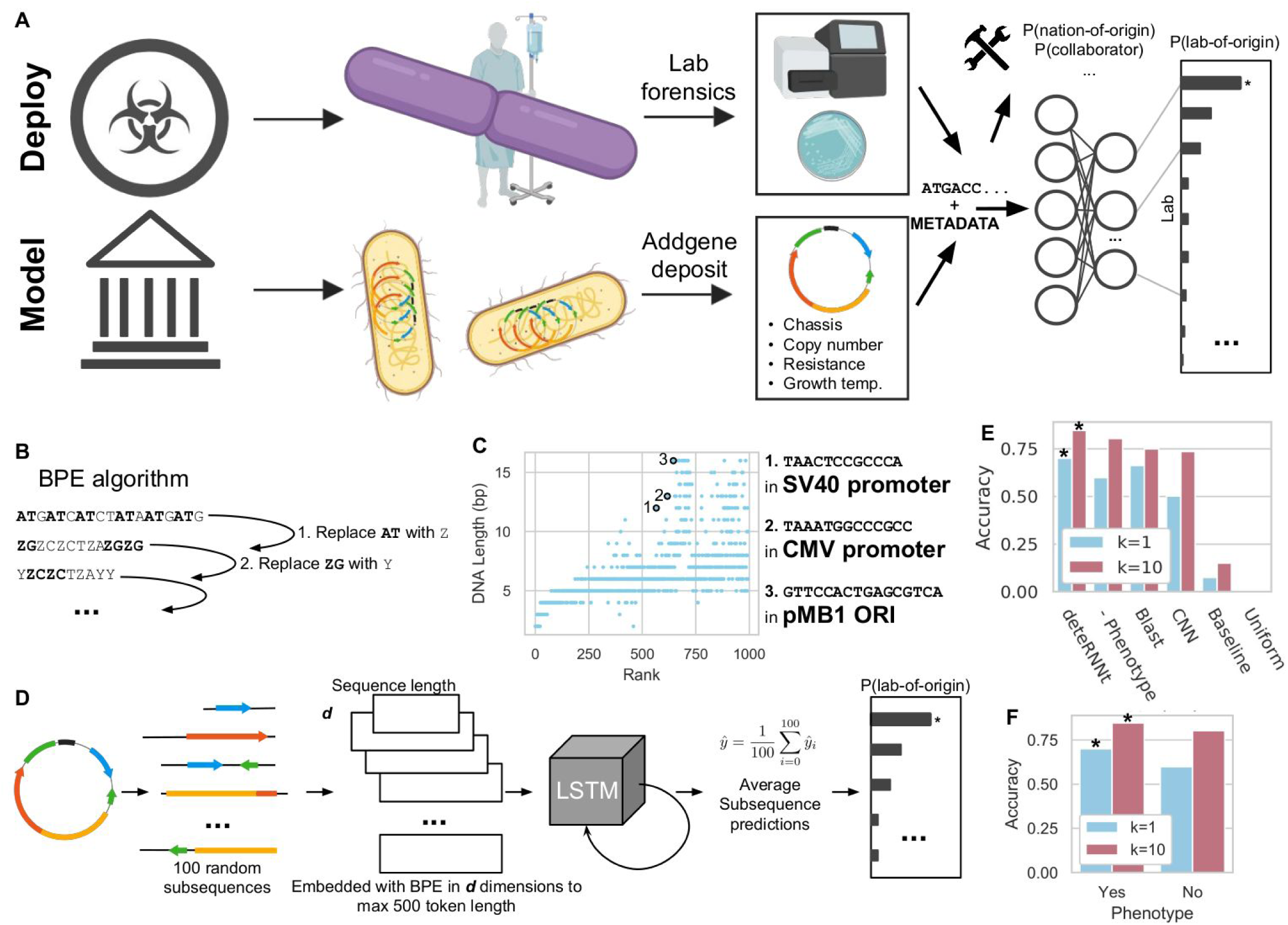
Deep learning on DNA motifs and minimal phenotype information enables state-of-the-art genetic engineering attribution. (**A**) The Addgene plasmid repository (bottom) provides a model through which to study the deployment scenario (top) for genetic engineering attribution. In the model scenario, research laboratories engineer organisms and share their genetic designs with the research community by depositing the DNA sequence and phenotypic metadata information to Addgene. In the corresponding deployment scenario, a genetically engineered organism of unknown origin is obtained, for example, from an environmental sample, lab accident, misuse incident, or case of disputed authorship. By characterizing this sample in the laboratory with sequencing and phenotype experiments, the investigator identifies the engineered sequence and phenotype information. In either case, the sequence and phenotype information are input to an attribution model which predicts the probability the organism originated from individuals connected to a set of known labs, enabling further conventional investigation. Above, the same information may be input to a wider toolkit of methods which provide actionable leads and characterization of the sample to support the investigator. (**B**) DNA motifs are inferred through the Byte Pair Encoding (BPE) algorithm^27^ which successively merges the most frequently occurring pairs of tokens to compress input sequences into a vocabulary larger than the traditional 4 DNA bases. Progressively, sequences become shorter and new motif tokens become longer. (**C**) BPE on the training set of Addgene plasmid sequences. The X axis shows the tokens rank-ordered by frequency in the sequence set (decreasing). The Y axis shows token length, in base pairs. Example tokens (bold, numbered) are linked to biologically meaningful sequence motifs. (**D**) The deteRNNt method takes 100 random subsequences from the plasmid encoded with BPE and embeds them into a continuous space via a learned word embedding^30^ matrix layer. These now variable length sequences are processed by a type of Recurrent Neural Network (RNN) called a Long-Short Term Memory (LSTM) network. We average the predictions from each subsequence to obtain a softmax probability that the plasmid originated in a given lab. (**E**) Top k prediction accuracy on the test set. Compared: deteRNNt, deteRNNt trained without phenotype, BLASTn, CNN deep learning state-of-the-art method^23^, a baseline guessing the most abundant labs from the training set, and guessing uniformly randomly (so low, cannot be seen).* indicates p < 10^−10^, by Welch’s t-test on n=30x 50% bootstrap replicates compared to BLAST. (**F**) Top k prediction accuracy on test set with and without phenotype information. * indicates p < 10^−10^, by Welch’s t-test on n=30x 50% bootstrap replicates compared to no phenotype.

## Results

DNA sequences are patterned with frequent recurring motifs of various lengths, like codons, regulatory regions, and conserved functional regions in proteins. Workhorse bioinformatic approaches like profile Hidden Markov Models^25^ and BLAST^26^ often rely on implicit or explicit recognition of local sequence motifs. Using an algorithm called Byte-Pair Encoding (BPE)^27^, which was originally designed to compress text by replacing the most frequent pairs of tokens with new symbols^28^, we inferred 1000 salient motifs directly from the Addgene primary DNA sequences (Fig. 1b, Methods) in a format suitable for deep learning. We found that the learned motifs appear to be biologically relevant, with the longer high-ranked motifs including fragments from promoters and plasmid origins of replication (ORIs) (Fig. 1c). Each of these inferred motifs is represented as a new token, so that a primary DNA sequence is translated into a motif token sequence that models can use to infer higher-level features.

Next, we prepared the Addgene dataset for training and model evaluation. The data were minimally cleaned to create a supervised multiclass classification task (Methods) in which a plasmid DNA sequence and 6 simple phenotype characteristics (Supplementary Table 1) are used to predict which of the 1,314 labs in the dataset deposited the sequence. This setup mirrors a scenario in which a biological sample is obtained, partially sequenced, and assayed to measure simple characteristics like growth temperature and antibiotic resistance (Supp. Table 1). The method can be trivially extended to incorporate any measurable phenotypic characteristic as long as a sufficiently large and representative dataset of measurements and lab of origin can be assembled, enabling integration with laboratory assays.

Before any analysis began, the data were split to withhold a ∼10% test set for evaluation (Methods). We further decreased the likelihood of overfitting by ensuring plasmids known to be derived from one another were not used in both training and evaluation (Methods).

### The deteRNNt model predicts lab-of-origin from DNA motif sequence and phenotypic metadata

For our model, we considered a family of Recurrent Neural Networks (RNNs) called Long-Short Term Memory (LSTM) networks, which process variable-length sequences one symbol at a time, recurrently, and are designed to facilitate the modelling of long-distance dependencies (Supplementary Fig. 1). We hypothesized that the intrinsically variable-length nature of RNNs would be well suited for biological sequence data and robust to data processing artifacts, and that attribution may rely on identifying patterns between distant elements, such as regulatory sequences and functional genes. We tokenized the sequences with variations of BPE, as described above, and searched over 250 configurations of architectures and hyperparameters (Supplementary Fig. 2-3, Methods).

The best-performing sequence-only model from this search (Supplementary Fig. 1) was then augmented with the phenotypic data (Supplementary Table 1) and fine-tuned (Methods). We elected to average the predictions of subsequences within each plasmid, as ensembling is widely recognized to reduce variance in machine learning models^29^. On an Nvidia K80 GPU, the resulting deteRNNt prediction pipeline (Fig. 1d) takes less than 10 seconds to produce a vector (summing to 1) of model confidences that the plasmid belongs to each lab. We compared Top-k lab-of-origin prediction accuracy on held-out test set sequences (Fig. 1e, Supplementary Table 2). Our model, with basic phenotype information, reaches 70.1% Top 1 accuracy. Our model additionally reached 84.7% Top 10 accuracy, a more relevant metric for narrowing down leads. This is a ∼1.7x reduction in the Top 10 error rate of a deep-learning Convolutional Neural Network (CNN) method previously considered the deep learning state of the art^23^ and similar reduction in BLAST Top 10 error. We note that BLAST cannot produce uncertainty estimates and has other drawbacks to deployment described elsewhere^23^. As expected, training without phenotype information reduced the Top k accuracy of our model (Fig. 1f).

### DeteRNNt is reasonably well calibrated and can be improved with temperature scaling

Having achieved lab-of-origin attribution accuracy of over 70%, we next considered the challenge posed to practical use by the black-box nature of deep learning. While powerful prediction tools, deep learning algorithms are widely recognized to be difficult to interpret^31,32^. Investigators of biological accidents or deliberate misuse will need to weigh the evidence from computational tools and other indicators, such as location, the incident’s consequences, and geopolitical context. Without this capability, computational forensic tools may not be practically useful. Consequently, as a starting point, we should demand that our models precisely represent prediction uncertainty: they must be well-calibrated. An attribution model is perfectly calibrated if, when it predicts that a plasmid belongs to Lab X with Y% confidence, it is correct Y% of the time. Performant deep learning models are not calibrated by default; indeed, many high-scoring models exhibit persistent overconfidence and miscalibration^33^.

Calibration can be measured empirically by taking the average model confidence within some range (e.g. 90-95%) and comparing it to the ground-truth accuracy of those predictions. Taking the average difference across all binned ranges from 0 to 100% is called the Expected Calibration Error (ECE). For situations where conservatism is warranted, the Maximum Calibration Error (MCE), which instead measures the *maximum* deviation between confidence and ground truth, can be used (Methods).

Following standard calibration analysis^33,34^, we find that our model is calibrated within 5% of the ground truth on average (ECE=4.7%, MCE=8.9%, Accuracy=70.1%, Fig. 2a). For comparison, the landmark image recognition model ResNet 110^35^ is within 16.53% of calibration on average on the CIFAR-100 benchmark dataset^33^. We next deployed a simple technique to improve the calibration of deteRNNt called temperature scaling^32^. After training, we learn a parameter called “temperature” which divides the unnormalized log probabilities (logits) in the softmax function by performing gradient descent on this scalar parameter to maximize the log probability of a held-out subset of data, our validation set. While more sophisticated techniques exist, temperature scaling provides a first-order correction to over or under-confidence by increasing or decreasing the Shannon entropy of predictions (Fig. 2a-b). After scaling, the re-calibrated model achieved a lower ECE of just 1.6% with a marginal decrease in accuracy (MCE=3.7%, Accuracy=69.3%, Fig. 2b, Methods). Using such calibrated predictions, investigators will be able to put more weight on a 95% prediction than a 5% prediction, choose not to act if the model is too uncertain, and have a basis for considering the relative importance of other evidence.

**Figure 2.**
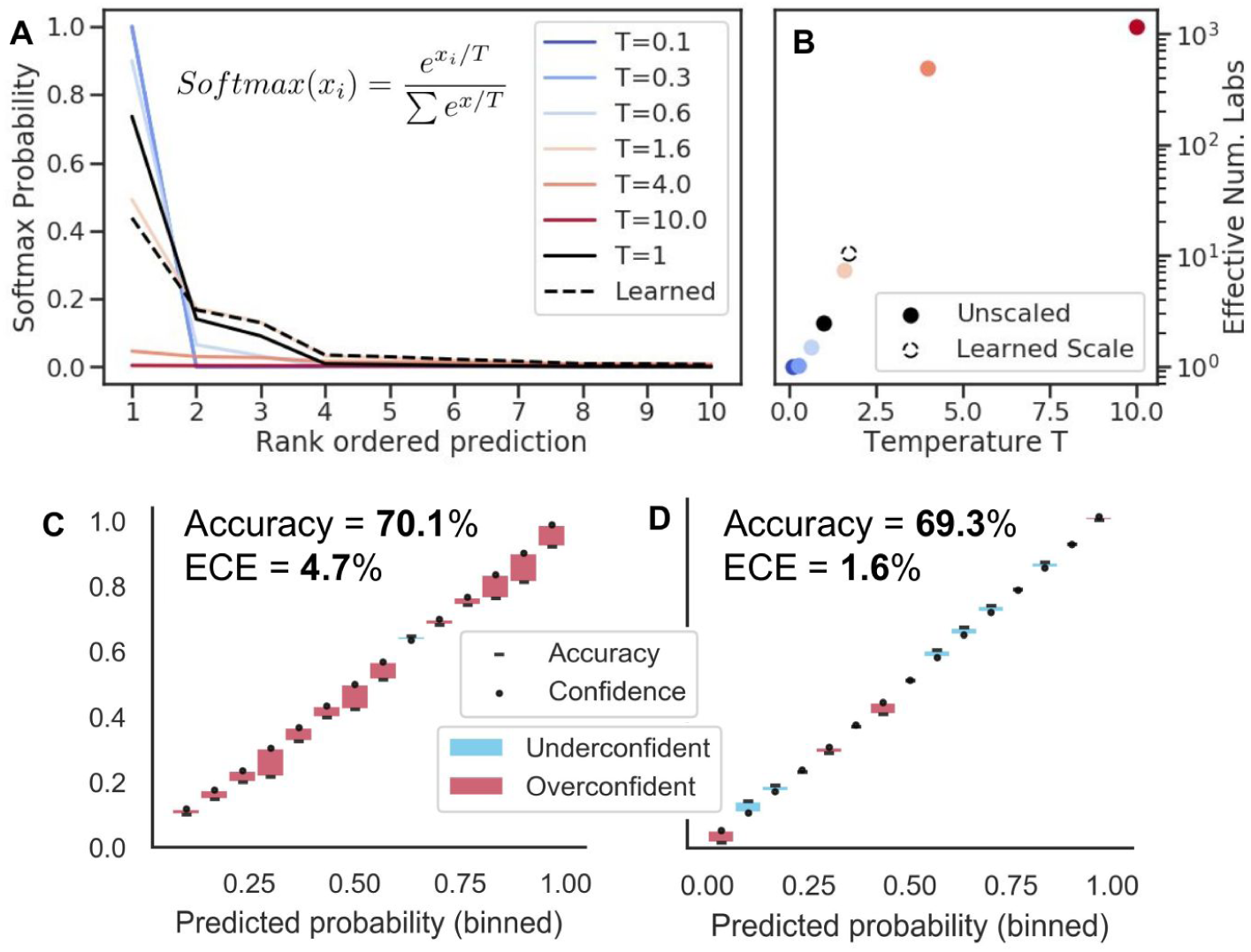
Model calibration and improvement with temperature scaling. (**A**) The effect of temperature scaling on the top 10 predictions of a randomly selected plasmid from the validation set’s predicted logits. The temperature parameter divides the class logits in the softmax, increasing (higher temperature) or decreasing (lower temperature) entropy of predictions. T=1 corresponds to default logits without temperature scaling. (**B**) Effective number of labs, exp(entropy), for the same random example as (A) as a function of temperature scaling parameter T with the unscaled default in black and the value of T learned by the temperature scaling calibration procedure shown in dotted outline. (**C**) Reliability diagram, pre-calibration. Every prediction in the test set is binned by predicted probability into 15 bins (x-axis). Plotted for each bin is the model’s predicted probability, or the confidence (dot) and the percentage of the time the model was correct within that bin, or the accuracy (dash). For a perfectly calibrated model, the accuracy and confidence would be equal within each bin. Instead, we see that the model is sometimes more confident than accurate (red regions) or more accurate than confident (blue regions). Overall prediction accuracy across all the bins and ECE are shown. (**D**) Same as (A) but after temperature scaling re-calibration.

### Expanding the toolkit: Predicting plasmid nation-of-origin with Random Forests

With a framework for calibration of attribution models that can in principle be applied to any deep learning classification algorithm, we next sought to expand the toolkit of genetic engineering attribution. While important, lab-of-origin prediction is only one component of the attribution problem. A fully developed framework would also include tools focused on laboratory attributes, which could narrow the number of labs to investigate, and the research dynamics by which designs collaboratively evolve, which could generate investigative leads. Furthermore, a suite of such tools may improve our understanding of the research process itself. Here, we begin lab attribute prediction with nation-of-origin prediction, which is particularly relevant for the Biological Weapons Convention (BWC). In terms of research dynamics, we begin by predicting the ancestry-descendent relationships of genetically engineered material.

For these analyses, we used a simple machine learning model called a Random Forest (RF)^36,37^, which ensembles decision trees, and encoded the DNA sequence of each plasmid with n-grams, which count the occurrence of n-mers in the primary sequence (Methods), instead of the more complex BPE-DNA motif encoding and highly parameterized deteRNNt model. We see this as a simple, but reasonably performant method for checking the feasibility of each task.

We inferred the nation-of-origin of a majority of the labs in the Addgene dataset through provided metadata and, in a few hundred cases, manual human annotation, without changing the boundaries of the train-validate-test split (Methods). There were 34 countries present after cleaning (Methods). We proposed a two-step nation-of-origin prediction process: 1) binary classification of a plasmid into Domestic v.s. International, to identify domestic incidents like local lab accidents or domestic bioterrorism; 2) multiclass classification to directly predict a plasmid’s nation of origin, excluding the domestic origin, to identify the source of an international accident or foreign bio-attack. We found the United States to make up over 50% of plasmid deposits in all the subsets (Fig. 4b) so we selected it as the domestic category for this analysis; in principle, any nation could play this role, and other training datasets would have different geographic emphasis. We trained an RF model, as described above, to perform binary U.S. vs International classification. As before, a ∼10% test set was withheld for evaluation (Methods). We found that RF performed comparably to BLAST (84.2% and 85.1%, respectively), but that both performed substantially better than guessing the most abundant class and uniformly choosing a class (Fig. 3a, RF ROC in Fig. 3b).

**Figure 3.**
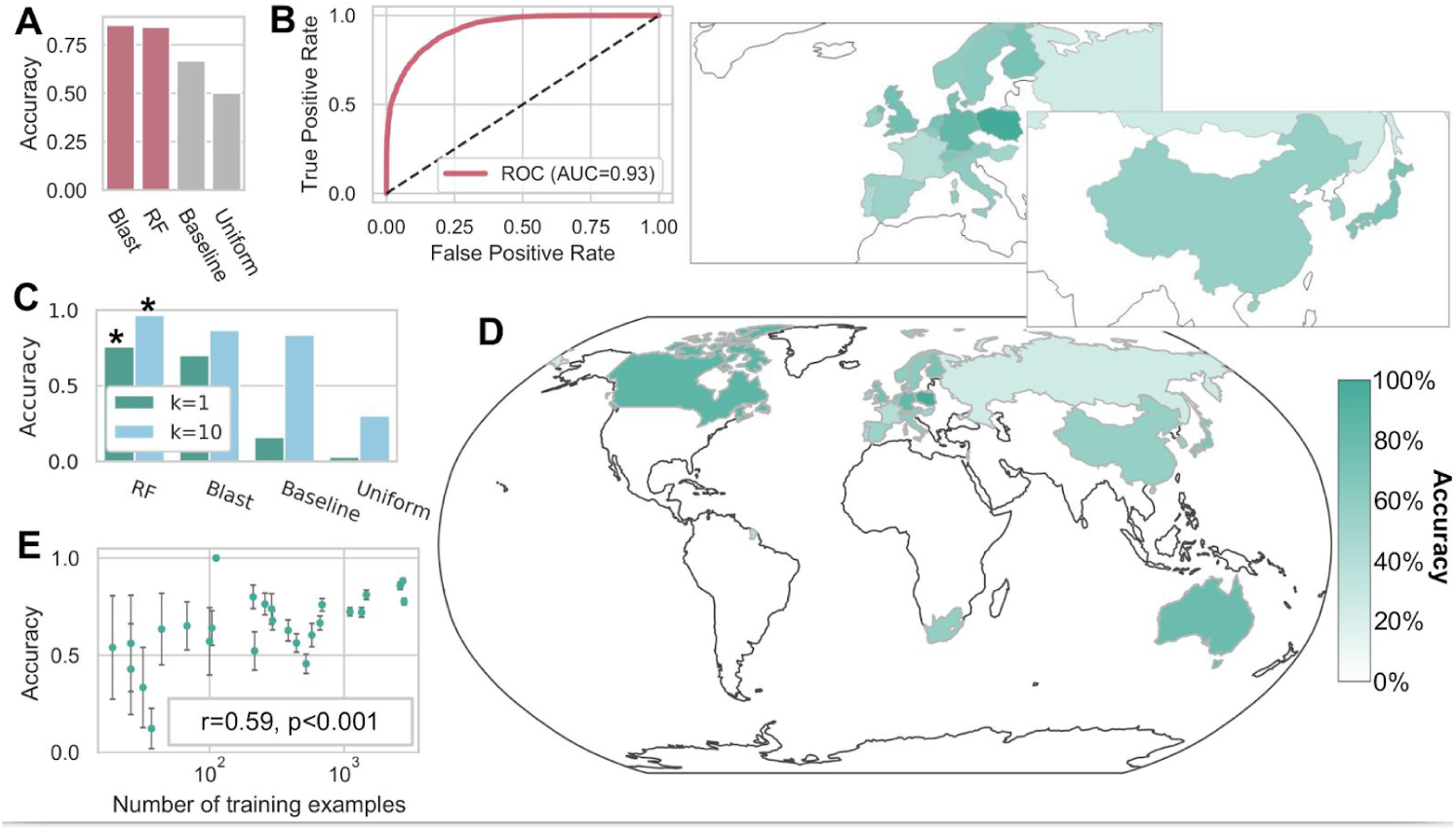
Simple machine learning methods can accurately attribute nation-of-origin of engineered DNA. (**A**) Domestic vs. International binary classification accuracy, with the United States defined as domestic of BLAST, Random Forests (RF), a baseline of predicting the most abundant class from the training set, and guessing uniformly randomly. Random Forests reach 84% accuracy. (**B**) Receiver Operating Characteristic (ROC) curve for the RF model, with Area Under the Curve (AUC) shown. (**C**) Top k test set accuracy of multi-class classification of nation-of-origin (excluding the United States, which is classified in (A)-(B)) for RF, BLAST and a baseline of predicting the most abundant class from the training set, and guessing uniformly randomly. With only 33 countries to choose between, guessing based on top 10 abundance reaches 83.8%. * indicates p < 10^−10^, by Welch’s t-test on 30x 50% bootstrap replicates compared to BLAST. (**D**) Test set accuracy of RF on multi-class classification of nation-of-origin, as in (C). Colored by prediction accuracy within each nation class. Enlarged Europe and East Asia shown above. (**E**) Nation-specific test set prediction accuracy correlates with the log10 of the number of training examples. Error bars represent standard deviations of prediction accuracies obtained by bootstrap-subsampling the test set examples used to evaluate the model.

**Figure 4.**
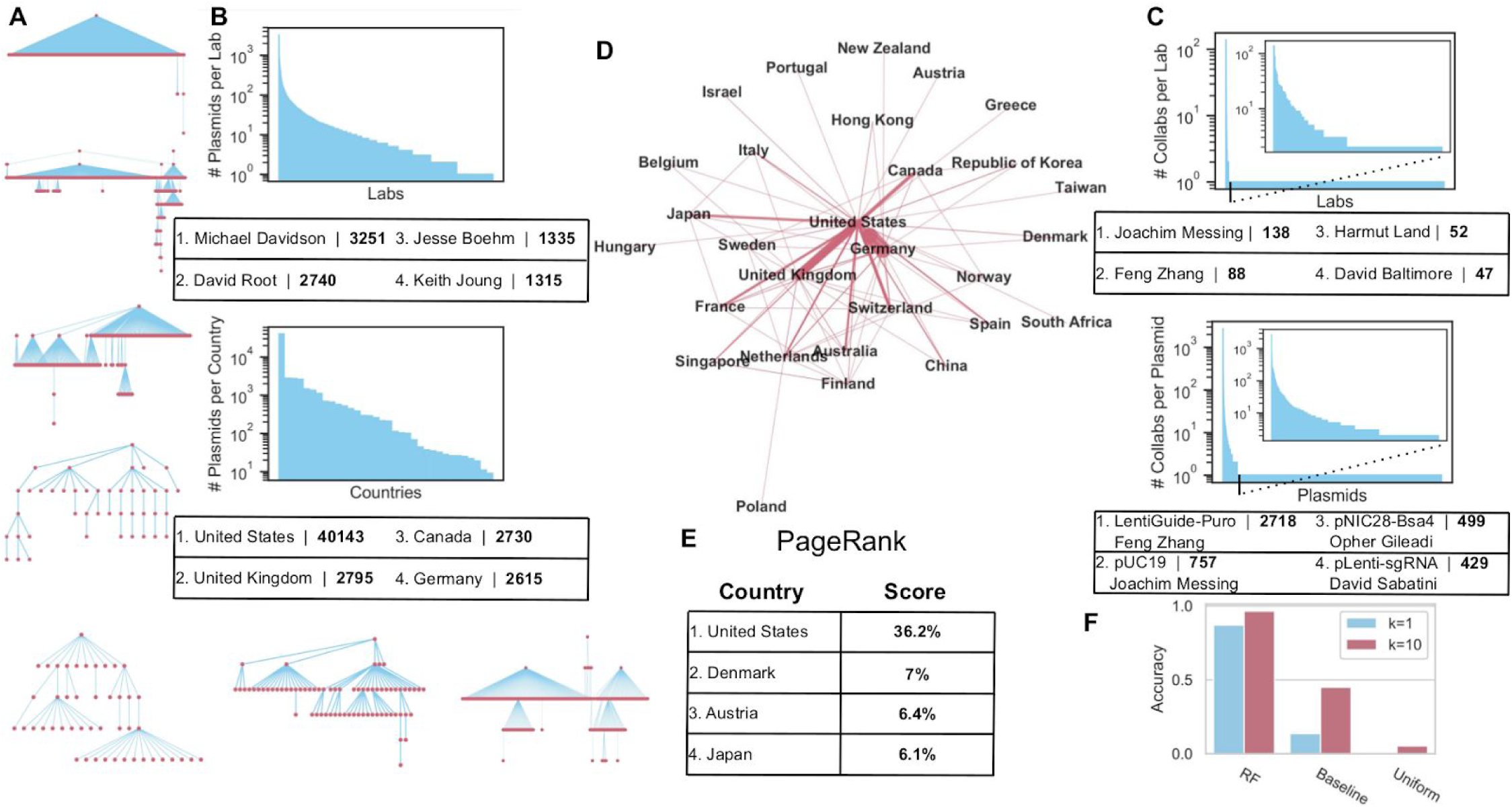
Lineage networks reveal skewed distributions in Addgene deposits and facilitate ancestor lab attribution with simple machine learning models. (**A**) Example ancestry-descendent lineages inferred from Addgene plasmid data. The largest lineage top, followed next down by the largest diameter, followed by a collection of examples from the more complex lineages. (**B**) Number of deposits. Above, number of plasmids deposited per lab, with ranked labs (descending) on the x-axis and log10 of the number of plasmids deposited on the y-axis. Top contributing labs in table. Below, number of deposits per country. (**C**) Number of ancestry-descendent connections. Above, number of ancestry-descendent linkages per lab, with most linked shown in the table as in (B). Below, number of linkages per plasmid. For each, subpanel shows the left side of the distribution by cutting out single linkages which constitute the right tail. (**D**) Lineage network between nations. Links indicate that at least one lab in a country has a plasmid derived from the other country or vice versa. Width of the link is proportional to the number of connected labs. (**E**) Top 4 PageRank scores from the lineage network from (D), represented as a directed weighted graph. The score is shown here as a percentage. (**F**) Top k accuracy predicting the ancestor lab: of a simple Random Forest (RF) model, the baseline of predicting the most abundant class(es) from the training set, and guessing uniformly randomly.

We proceeded to nation-of-origin prediction with the remaining international country assignments. A multi-class RF model was trained on the training set. Prediction accuracies within each country varied substantially (Fig. 3d) but overall RF accuracy was 75.8% with Top 10 accuracy reaching 96.7% (Fig. 3c, Supplementary Table 3). If asked to predict the top 3 most likely countries, which could be more useful for investigators, we find 87.7% Top 3 accuracy for RF compared to 47.0% for guessing the abundant classes. As shown in Fig. 3e, classification performance is degraded by low sample size for many countries in the dataset, explaining some of the variability in Fig. 3d. We therefore expect performance to improve and variability to be reduced as more researchers in these countries publish genetic designs. Together, these results suggest that it may be possible to model genetic engineering provenance at a coarse-grained level, which should be helpful in cases where a lab has never been seen before. However, the dataset used here has substantial geographic bias due to Addgene’s location in North America and difference in data and materials sharing practices as a function of geography. Additionally, the accuracy of our simple model could be substantially improved. Even with those considerations addressed, improved nation-of-origin models, as with the other methods in this paper, should be one part of an integrated toolkit that assists human decision makers rather than assigning origin autonomously.

### Expanding the toolkit: Inferring collaboration networks by predicting ancestor lab-of-origin

We continued to expand the toolkit of attribution tools by evaluating the feasibility of predicting plasmid research dynamics in the form of ancestry-descendent relationships. These might help elucidate connections between research projects, improve credit assignment and provide evidence in cases when the designer lab has not been observed in the data but may have collaborated with a known lab. Ancestry relationships may be caused by deliberate collaboration between one lab sharing an “ancestor” plasmid with another lab who creates a derivative, or by incidental reuse of genetic components from published work. In our usage, these ancestry-descendent relationships form “lineages” of related plasmids.

We inferred plasmid ancestor-descendent linkages from the relevant fields in Addgene which acknowledge sequence contributions from other plasmids (Methods). By following these linkages, we identified 1223 internally connected lineage networks of varying sizes (example networks in Fig. 4a). Analysis of the networks revealed that the number of plasmids deposited by lab and country (Fig. 4b), number of lab-to-lab connections, and number of connections per plasmid (Fig. 4c) exhibited extremely skewed, power-law like distributions (Methods). These patterns reflect some combinations of the trends in who deposits to Addgene and the underlying collaboration process.

Next, we constructed an international lineage network: a weighted directed graph of ancestry-descendent connections between labs in different countries, with weights proportional to the number of unique connections from lab to lab (Fig. 4d, Methods). We used the Google PageRank algorithm, which scores the importance of each country in the network by measuring the percentage of time spent in each country if one followed the links in the network randomly (Fig. 4e, Methods). As expected from the geographic centrality of the United States in this dataset, it scores by far the highest at 36.2%. Unexpectedly, Denmark, Austria, and Japan followed as the next most important nodes despite not being in the top 4 countries in terms of raw Addgene contributions.

Finally, we labelled each plasmid with the lab of its most recent ancestor, e.g. the lab which contributed part of the source sequence of the plasmid. We call this ancestor lab attribution, to distinguish that it occurs one step above lab-of-origin attribution in the lineage network. We re-split the data to ensure representation of each ancestor lab in the training and test sets (Methods). On the held-out test set, we find Top 1 accuracy of 87.0% and Top 10 accuracy of 96.5%, compared to guessing the most frequent class(es) at 13.6% (k=1) and 45.0% (k=10) respectively (Fig. 4f, Supplementary Table 4). We note that this high accuracy compared to standard lab-of-origin prediction is likely explained by having only 188 ancestor labs to guess between (compared to 1,314 previously), and is limited by the lineage structure we were able to infer from Addgene, which may be particular to this unique resource instead of representative of broader trends in collaboration.

### Exploring the deteRNNt model

A better understanding of the predictions of and features learned by neural networks for attribution is desirable. However, while model calibration can be a universal criteria for any black-box model, it is notoriously challenging (and often model specific) to visualize and understand the high-dimensional non-linear function of the input data which a deep neural network represents^30,31^.

We began by visualizing the learned feature representation of the model. We took the model’s 1000-dimensional hidden activations and projected them into 2 dimensions using a t-distributed stochastic neighbor embedding^39^ (tSNE) (Methods). We saw many large, well-separated clusters of plasmids that are assigned high probability by the model (Fig. 5a). This is in line with the expectation that a model can classify a group of plasmids more accurately if they are more separable in the hidden space.

**Figure 5.**
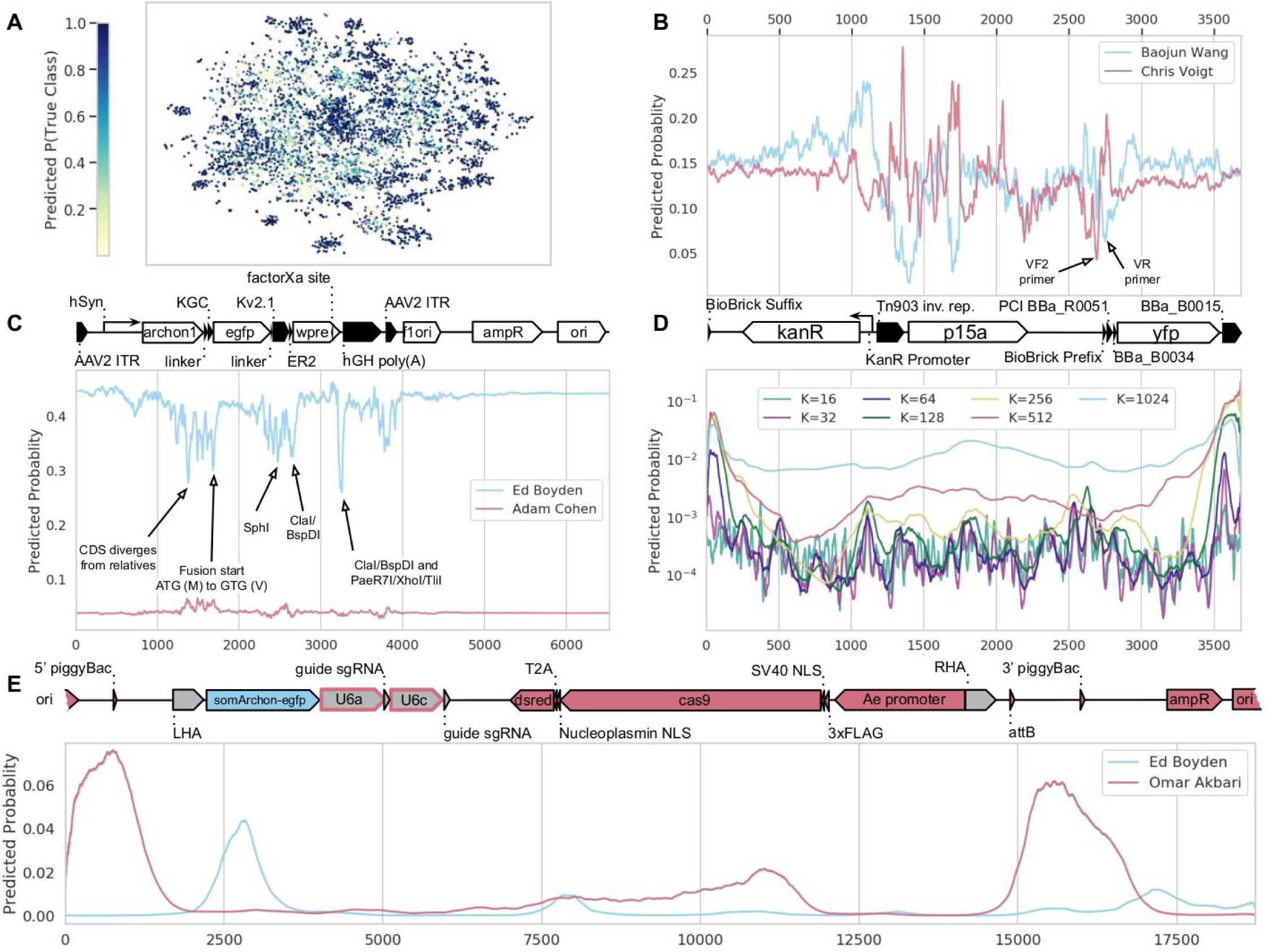
Interpreting the deteRNNt model. (**A**) 2-dimensional tSNE of the pre-logit hidden layer of deteRNNt for the validation set. Colored by probability assigned to the true lab. An interactive 3-d visualization is available at http://papers.altlabs.tech/visualize-hiddens.html. (**B**) Scanning mask of 10 N’s across linear sequence (x-axis) of Chris Voigt lab plasmid pCI-YFP, (Genbank JQ394803.1) on the brink of predicting Baojun Wang Lab vs. Voigt Lab. At each x position is shown the probabilities given by softmax on the mean of the logits from all the sequences which include that position masked with N. Plasmid schematic is shown below. (**C**) The scanning 10-N mask analysis from (B) applied to Ed Boyden Lab Plasmid pAAV-Syn-SomArchon (Addgene #126941). The second-most likely lab predicted by deteRNNt on this plasmid is shown in red, with plasmid schematic above. (**D**) Predicting lab-of-origin from subsequences of varying lengths K scanned across the linear sequence. At each x the probabilities are given by softmax of the mean of logits from subsequences which include that position in the window. Color indicates subsequence K-mer length. Linear sequence position is given by the x-axis and shared with (B) with plasmid schematic above. (**E**) Custom designed gene-drive plasmid derived from an Omar Akbari *Aedes aegypti* germline Cas9 gene drive backbone (AAEL010097-Cas9, Addgene #100707) carrying a payload of Cas9-dsRed, a guide cassette, and SomArchon from the Boyden plasmid pAAV-Syn-SomArchon from (C), with scanning subsequence window as in (D) and K=1024. Predicted probabilities for Omar Akbari (red) and Boyden (blue) are shown, along with the correspondingly colored plasmid schematic. All unlabelled regions are from the Akbari vector. The right and left homology arms shown in grey were introduced by our design and come from neither lab’s material, and the U6a and U6c were from Akbari lab material (Addgene #117221 and #117223, respectively) but introduced into the backbone by our design (grey outlined in red).

We next examined three case studies to inspect the contribution of DNA sequence features to deteRNNt predictions. These should be taken as suggestive, given the anecdotal nature of analyzing an example plasmid. We first looked at a plasmid from Chris Voigt’s lab, pCI-YFP (Genbank JQ394803.1) which is not in Addgene and has been used for this purpose in prior work^23^. pCI-YFP is notable because it is composed of very widespread and frequently used components (Fig. 5b below). We found that sequence-based deteRNNt is uncertain whether this plasmid was designed by the Voigt Lab or by Baojun Wang’s Lab (around 25% probability to each). Notably, the Wang lab does substantial work in genetic circuit design and other research areas which overlap with the Voigt lab. We adapted a method described previously^23^ to perform an ablation analysis, where the true DNA sequence within a window is replaced by the unknown symbol N. We scan a window of 10 N’s across the length of the sequence and predict the lab of origin probabilities for each.

We visualized what sequences were critical for predicting the Voigt lab (drop in Voigt probability when ablated) and which were “distracting” the model into predicting the Wang label (rise in Voigt probability when ablated) (Fig 5. b). Among the sequence features we identified, we found a sequence in a Tn903 inverted repeat (upstream of the p15a origin of replication) which, when deleted, causes a spike in Voigt Lab probability, and also found two unlabelled primer sites, VF2 and VR which seem to be important for predicting Voigt and Wang, respectively.

Given the age of pCI-YFP (published in 2012) we were concerned that it might not be the most representative case study because 1) genetic engineering is a rapidly evolving field and more recent techniques and research objectives could likely have shifted the data distribution since this plasmid’s design and publication, and 2) older plasmids are more likely to have their subcomponents used in other plasmids, especially within the same lab where material transfer is instantaneous.

We identified the plasmid pAAV-Syn-SomArchon (Addgene #126941) from Ed Boyden’s lab for an additional case study. This plasmid was also outside our dataset because it was published^40^ after our data was downloaded. Beyond representing a case of generalizing forward in time and to an unpublished research project, this plasmid also contains the coding sequence for SomArchon, which is the product of a screening-based protein engineering effort. We note that despite being a design newer than our data, this plasmid builds on prior work, and we expect some microbial opsins and elements of the backbone to be present elsewhere in Addgene.

Given the entire sequence, sequence-based deteRNNt predicts the plasmid belongs to the Boyden Lab as its top choice at 44%, followed by the “Unknown Engineered”’ category at 10%, followed by the Adam Cohen lab at 4%, which has also published on microbial-opsin based voltage indicators^41,42^. Performing the same analysis as above, we examined the predictions while scanning 10 Ns along the sequence (Fig. 5c). Interestingly, there were many restriction sites that when deleted dropped the Boyden probability. Restriction sites choices can often be a hallmark of a research group. In addition, the choice of what to do to the ATG start codon in a fusion protein (in this case the eGFP fused to ArchonI) can differ between labs. One could leave it be, delete it, or mutate it, as the Boyden group chose to do in this case (ATG to GTG, M to V), which our model appears to identify as a critical feature (drop in Boyden probability of ∼10% when deleted) (Fig. 5c).

Unlike architectures which assume a fixed-length sequence, deteRNNt can accept any length sequence as input. We elected to use this capability to examine the marginal contribution of K-mer subsequences of various length to the overall prediction. Unlike the analysis above, this tests what the model believes about a fragment of the plasmid in isolation. We selected a sliding window of subsequence of length K across pCI-YFP and visualized the positional average Voigt lab predicted probability (Fig. 5d). We see that in general subsequence predictions produce probabilities substantially lower than the full-sequence ∼25%, with higher-frequency features showing much lower predicted probability. K-mers containing BBa_B0015 on the right hand terminus of the sequence have a surprisingly high marginal probability predicting Voigt, around 10% for K=256, while the ablation analysis does not identify this as a critical region.

We were curious to what degree the predictions of deteRNNt on these two case studies were dependent on having the highly discriminative motifs, like restriction sites and codon selection noted above, co-occur with backbone elements which are incidental to the functioning of the system. In practice, the backbone plasmid may substantially change as part of a new project within a lab, or an actor who is not inside addgene may combine elements from existing plasmids for a new purpose. While we can’t expect a multi-class classification model to identify a never-before-seen lab, we might hope that it could identify which components of a plasmid were likely derived from, or designed by, a given lab. If this subcomponent attribution could be achieved, it might provide another useful tool for understanding the origin of an unknown genetic design.

To test this we designed a novel gene drive plasmid for use in *Aedes aegypti* by starting from a germline Cas9 backbone designed in the Omar Akbari lab (AAEL010097-Cas9, Addgene #100707). In our design, the original Cas9 is fused with dsRed from elsewhere in the plasmid, and this backbone was modified to introduce homology arms and include the payload of a guide RNA cassette and the SomArchon CDS from the Boyden Lab’s pAAV-Syn-SomArchon examined above (Methods). As we might expect, the model is very uncertain about this hybrid plasmid, and assigns ∼35% probability to it being from the “Unknown Engineered” class. Among the true lab classes it assigns the Akbari lab highest probability at ∼3%, but only gives the Boyden lab .3%. When we apply the scanning K-mer analysis from Fig. 5d, with K=1024, we find that the model peaks its Akbari lab predictions on backbone elements around two piggyBac sites, and furthermore the Boyden lab-derived SomArchon payload has a peak of predicted Boyden probability, indicating that the model could identify this functional payload even absent clues from the pAAV-Syn-SomArchon backbone. While these analyses should be taken as anecdotes as with any case-study, we note that future work could explicitly model the attribution problem as predicting the lab-of-origin of every position in a sequence.

## Discussion

Together, these results suggest a practical and accurate toolkit for genetic engineering forensics is within reach. We achieve 70% accuracy on lab-of-origin prediction, a problem previously thought challenging if not intractable. Our work has the advantages of using biologically motivated, motif-based sequence models, and leveraging phenotype information to both improve accuracy and interface with laboratory infrastructure. Furthermore, with model calibration we provide the first framework for weighing the predictions of attribution forensics models against other evidence. Finally, we establish new attribution tasks — nation of origin and ancestor lab prediction— which promise to aid in bioweapons deterrence and open the possibility of more creative attribution technologies. While we focus here on security, attribution has wide implications. For example, attribution could promote better lab safety by tracing accidental release^43^. Additionally, we discovered an interesting, albeit anecdotal, power-law like skew in Addgene plasmid deposits, which may reflect the particular dynamics of this unique resource, but is also consonant with prior work on scale-free patterns of scientific influence^44–46^. We believe that a deeper understanding of the biotechnology enterprise will continue to be a corollary of attribution research, and perhaps more importantly, that computational characterization tools, like ancestor lab prediction, will directly promote openness in science, for example by increasing transparency in the acknowledgement of sequence contributions from other labs, and establishing a mechanism by which community stakeholders and policy makers can probe the research process.

Our analysis has limitations. Machine learning depends on high-quality datasets and the data required to train attribution models is both diverse and scenario dependent. That said, we believe it is the responsibility of the biotechnology community to develop forensic attribution techniques proactively, and while the Addgene attribution data provides a reasonable model scenario, future work should look to build larger and more balanced datasets, validate algorithms on other categories of engineered sequences like whole viral and bacterial genomes, and analyze their robustness to both obfuscation efforts and dataset shifts, especially considering new methods for robust calibration^47^. We further note that 70% lab-of-origin attribution accuracy is not conclusive on its own, and any real investigation with tools at this level of accuracy would do well to almost entirely rely on human expertise. However, combined with traditional investigation, microbial forensics^3,4^, isotopic ratio analysis^48–50^, evolutionary tracing^51^, sequence watermarking^51,52^, and a collection of machine learning tools targeting nation states, ancestry lineages, and other angles, our results suggest that a powerful integrated approach can, with more development, amplify human expertise with practically-grounded forensic algorithms. In the meantime, developing automated attribution methods will help scale efforts to understand, characterize and study the rapidly expanding footprint of biotechnology on society, and may in doing so promote increased transparency and accountability to the communities affected by this work.

We see our results as the first step towards this integrated approach, yet more work is needed. The bioengineering, deep learning, and policy communities will need to creatively address multidisciplinary problems within genetic engineering attribution. We are hopeful that the gap between these fields can be closed so that tools from deep learning and synthetic biology are proactively aimed at essential problems of responsible innovation.

## Acknowledgments

We thank Grigory Khimulya, Gregory Lewis, Michael Montague, Andrew Snyder-Beattie, Surojit Biswas, Mohammed AlQuraishi, Gregory Koblentz, and Gabriella Deich for valuable feedback and discussion. We also thank Christopher Voigt and Alec Nielsen for their critical commentary. We thank Addgene, particularly Jason Niehaus and Joanne Kamens, for data access and thoughtful feedback.

## Author Contributions

E.C.A. conceived the study and designed the analyses. M.T. processed and cleaned the data. T.K. split the data. M.T. managed data and software for part of Random Forest and network analysis. A.B.L. managed data and software for part of nation-state and ancestor analysis. J.S. managed part of the CNN baseline. E.C.A. managed data and software for all other analyses. R.E. and S.E.VS. designed the gene-drive plasmid and performed sequence analysis to interpret model predictions. G.M.C. and K.M.E. supervised the project. E.C.A. wrote the manuscript with help from all authors.

## Methods

### Graphing and visualizations

All graphs and plots were created in python using a combination of the seaborn (https://seaborn.pydata.org/), matplotlib (https://matplotlib.org/3.1.0/index.html), plotly (for geographic visualization and interactive 3d hidden state visualization: https://plot.ly/) and networkx (https://networkx.github.io/).

### Processing the Addgene Dataset

The dataset of deposited plasmid sequences and phenotype information was used with permission from Addgene, Inc.

Scientists using Addgene may upload full or partial sequences along with metadata such as growth temperature, antibiotic resistance, vector backbone, vector manufacturer, host organism, and more. For quality control, Addgene sequences portions of deposited plasmids, and in some cases sequences entire constructs. As such, plasmid entries featured sequences categorized as addgene-full (22,937), depositor-full (32,669), addgene-partial (56,978), depositor-partial (25,654). When more than one category was listed we prioritized plasmids in the order listed above. When there was more than one entry for a category, the longest sequence was chosen. If more than one partial sequence was present, we concatenated them into a single sequence. The final numbers by sequence category were addgene-full (22937), depositor-full (28465), addgene-partial (27185), and depositor-partial (3247). Plasmids were dropped if they did not have any registered sequences. Any Us were changed to Ts and letters other than A, T, G, C, or N were changed to Ns. The resulting dataset contained 81834 plasmids, from 3751 labs.

The raw dataset contains as many as 18 metadata fields from Addgene. In the final dataset we kept only host organism species, growth temperature, bacterial resistance, selectable markers, growth strain, and copy number. These fields were selected because they are phenotypic characteristics we expect to be easy to measure in the scenario considered here, where a sample of the organism is available for sequencing and wet-lab experimentation. For more information on what these phenotyping assays might look like, see Supplementary Table 1.

Metadata fields on Addgene are not standardized and have many irregularities as a result. To deal with the lack of standardization, our high-level approach was to default to conservatism in assigning a given plasmid some phenotype label. We used an “other” category for each field to avoid noisy or infrequent labels, in particular we assigned any label that made up less than 1% of all labels in some field to “other”. Additionally, some of the fields had multiple labels e.g. the species field may list multiple host organisms: “H. sapiens (human), M. musculus (mouse), R. norvegicus (rat)”. These fields were one-hot encoded, allowing multiple columns to take on positive values if multiple labels were present. If the field contained no high frequency labels (more than 1%) and was not missing, the “other” column was set to 1. A small number of plasmids (117) had sequences but no metadata. While far from perfect, our choice to use the default category of “other” to avoid introducing noisy information should prevent spurious features from being introduced by including phenotypic metadata. Given the performance boost achieved by including even this very minimal phenotype information, we are enthusiastic about future efforts to collate more standardized, expressive and descriptive phenotype information.

### Inferring Plasmid Lineage Networks

Many plasmids in the Addgene database reference other plasmids used in their construction. Within the metadata of each plasmid, we searched for references to other sequences in the Addgene repository, either by name or by their unique Addgene identifier. Plasmid names were unique except for 1519 plasmids that had names associated with more than one Addgene ID (331 of these also had duplicate sequences). However, none of these plasmids with duplicate names were referenced by name by some other plasmid. We considered a plasmid reference in one of the following metadata fields to be a valid reference: *A portion of this plasmid was derived from a plasmid made by, Vector backbone, Backbone manufacturer*, and *Modifications to backbone*. Self-references were not counted, and in the rare case where two plasmids referred to each other, the descendent/ancestor relationship was picked at random.

We were interested in discovering networks of plasmids with shared ancestors— collectively we may call this subset of plasmids a lineage network. The problem of assigning plasmids to their associated network reduces to the problem of finding a node’s connected component from an adjacency list of a directed, potentially cyclic, graph. The algorithm proceeded iteratively: in each round we picked some unvisited node. We then performed breadth-first search (plasmids were allowed to have multiple ancestors) and assigned all nodes visited in that round to a lineage network. In the case where a visited node pointed to a node that was already a member of some network, the two networks were merged. Keeping track of nodes visited in each round prevented the formation of cycles. We verified this result by reversing our adjacency list and running the same algorithm, equivalent to traversing by descendents instead of ancestors.

### Train-Test-Validation Split

To rigorously evaluate the performance of a predictive algorithm, strong boundaries between the datasets used for training and evaluation are needed to prevent overfitting. We follow best practices by pre-splitting the lab-of-origin Addgene data into an 80% training set, ∼10% validation set for model selection, and ∼10% test set held out for final model evaluation. In our vocabulary, the training set can be used for any optimization, including fitting an arbitrarily complex model. The validation set may only be used to measure the performance of an already trained model, e.g. to select architecture or hyperparameters; no direct training. Finally, the test set may only be used after analysis is completed, the architecture and hyperparameters are finalized, as a measure of generalization performance. We addressed two additional considerations with the split:

1. A large number of labs only deposited one or a few sequences. This is insufficient data to either train a model to predict that lab class or reasonably measure generalization error.
2. Like many biological sequence datasets, the Addgene data are not independent and identically distributed because many plasmids are derived from others in the dataset, potentially creating biased accuracy measures due to overperformance on related plasmids used for both training and evaluation.

To handle the first, we choose to pool plasmids with fewer than 10 examples into an auxiliary category called “Unknown Engineered”, and additionally stratify the split to ensure that every lab has at least 3 plasmids in the test set.

For the second, we inferred lineage networks (see above). We stratified the split such that multi-lab networks were not split into multiple sets. In other words, each lineage was assigned either to the training, validation, or test set as a group, not divided between them.

We used the GroupShuffleSplit function in python with sklearn (http://scikit-learn.org/stable/) to randomly split given these constraints. The final split had 63,017 training points, 7,466 validation points, and 11,351 test points. The larger test set is a direct result of enforcing ⅓ of rare lab’s plasmids are split there for generalization measurement. We note that, because model selection is occurring on the validation set which has no representation of a number of rare labs, there is a built-in distributional shift that makes generalizing to our test set particularly challenging. However, we believe that this is appropriate to the problem setting— attribution algorithms should be penalized if they cannot detect rare labs, because in a deployment scenario the responsible lab may be unexpected. A visualization of this phenomena, and the lab distribution after splitting can be found in Supplementary Fig. 4.

We confirmed that this cleaning and splitting procedure did not dramatically change the difficulty of the task from prior work by reproducing the model architecture, hyperparameters, and training procedure of a model with known performance on a published dataset (see Baselines in Methods)^23^.

### Byte Pair Encoding

The sequences from the training set were formatted as a newline separated file for Byte Pair Encoding inference. Inference was performed in python on Amazon Web Services (AWS) with the sentencepiece package (https://github.com/google/sentencepiece) using both the BPE and Unigram^53^ algorithms, with vocabulary sizes in [100, 1000, 5000, 10000], no start or end token, and full representation of all characters. The resulting model and vocabulary files were saved for model training, which used sentencepiece to tokenize batches of sequence on the fly during training. For the Unigram model, which is probabilistic, we sampled from the top 10 most likely sequence configurations.

For the visualization and interpretation of the 1000-token BPE vocabulary ultimately selected by our search algorithm, we took the vocabulary produced by sentencepiece, which has a list of tokens in order of merging (which is based on their frequency), and plotted this ordering vs. the length of the detected motif. We selected 3 example points visually for length at a given ranking. These sequence motifs were interrogated with BLAST^26^ with the NCBI web tool, and additionally with BLAST tool on the iGEM Registry of Standard Biological Parts (http://parts.igem.org/sequencing/index.cgi). For each motif, a collection of results were compared with their plasmid maps to place the motif sequences within a plasmid component. We found, as numbered in the figure, motif 1 repeated twice in the SV40 promoter, motif 2 repeated twice in the CMV promoter, and motif 3 occuring slightly downstream of the pMB1 origin of replication.

### Training deep Recurrent Neural Networks

We consider a family of models based on the LSTM recurrent neural network and a DNA motif-base encoding. We formulated an architecture and hyperparameter search space based on prior experience with these models. In particular, we searched over categorical options of learning rate, batch size, bidirectionality, LSTM hidden size, LSTM number of layers, number of fully connected layers, extent of dropout, class of activations, maximum length of the input sequence and word embedding dimension. We further searched over parameters of the motif-based encoding, including whether it was Unigram or BPE based and the vocabulary size. Configurations from this categorical search space were sampled and evaluated by the Asynchronous Hyperband^54^ algorithm, which evaluates a population of configurations in tandem and halts poor performing models periodically. Thus, computational resources are more efficiently allocated to the better performing models at each step in training. Our LSTM architecture was optimized with Adam using categorical cross-entropy loss in PyTorch (https://pytorch.org/) and hyperparameter tuning with Asynchronous Hyperband used ray tune (https://ray.readthedocs.io/en/latest/tune.html).

In early exploratory experiments, we found that including metadata in the initial training process caused a rapid increase but quick plateau of the Hyperband population. We noticed that these models usually had small LSTM components, suggesting that they were ignoring sequence information. This led us to hypothesize that adding metadata early in training led to an attractive local minima for the tuning process which neglected sequence, exploiting the fact that Hyperband penalizes slow-improving algorithms.

We therefore adopted a progressive training policy as follows. First, 250 configurations of the search space described above were evaluated with ray tune (Supplementary Fig. 2-3) over the course of ∼1 week on an AWS p2.8xlarge machine with K80 GPUs. The best performing model was selected for a stable and steadily decreasing loss curve (Supplementary Fig. 3) after 300 Hyperband steps, each of 300 weight updates (∼90,000 updates total). This model configuration was saved and trained from scratch to ∼200,000 weight updates, selected based on an early stopping heuristic on the validation loss. Next, this model was truncated to the pre-logit layer, and metadata was concatenated with the output of this sequence only model (Supplementary Fig. 1), followed by one hidden layer and the logit layer. This was further trained, but with the LSTM sequence model frozen until validation loss plateaued. Finally, the full model was jointly trained until validation loss plateaued. The effect of this approach was to prevent the model converging on a metadata-focused local optimum without overfitting on the training set (which was facilitated by training only components of the model at a time).

After fully training and finalizing results using our original random split (see above), 3 additional random splits were performed and training was repeated as before using different random seeds but the same hyperparameters as were found in the first hyperband search. We found that even without hyperparameter tuning on each newly split dataset, the results for the full and sequence-only deteRNNt models were consistent with our earlier results (Supplementary Fig. 5).

### Calibration analysis

We followed the methodology of Guo et al (2017)^33^. Prediction on the test set were binned into 15 bins, *B*_*m*_. The difference between the confidence of a model and the accuracy of the resulting model’s predictions, termed calibration can be measured by two metrics, Expected Calibration Error (ECE) measuring the average difference between prediction confidence and ground truth accuracy, and Maximum Calibration Error (MCE) measuring the maximum thereof. They are defined below (n is the number of samples).

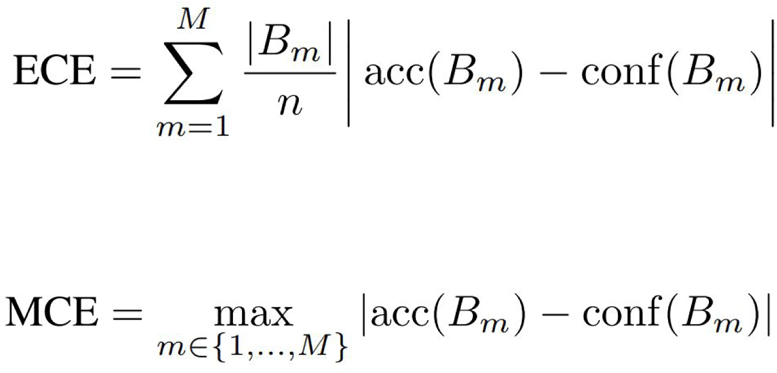

### Temperature Scaling

Temperature scaling adjusts the logits (pre-softmax output) of a multiclass classifier) by dividing them by a single scalar value called the temperature. For a categorical prediction *q*, logits **z** and temperature T, we have:

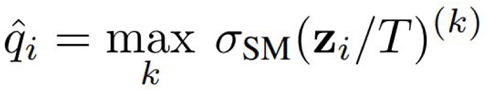

The temperature is learned on the validation set after the model is fully trained. We used PyTorch and gradient descent with the Adam optimizer to fit the temperature value. Subsequently, all of the logits predicted for the test set by the original model were divided by the temperature and softmaxed to get confidences as shown in Fig. 2 (right). Because the maximum of the softmaxed vector is mathematically equivalent to the maximum of a softmax on scaled logits, we concluded that the slight difference in accuracy of the calibrated model was due to floating-point errors.

### Random Forest models

We used scikit-learn package (https://scikit-learn.org/stable/index.html) implementation of Random Forest Regressor. Unless otherwise specified, we used 1000 estimators with 0.5 as the maximum proportion of features to look at while searching for the best split and with class weights inversely proportional to class frequency.

For Random Forest analysis we represented the sequences as frequencies of 1,2,3 and 4-grams. We constructed the ngram vocabulary using the training set, and then only used the frequencies of ngrams included in the vocabulary to construct features (ngram frequencies) for the validation and test sets. Where specified, we concatenated one-hot-encoded metadata (phenotypic information) to these ngram frequencies. We transformed the ngrams features using TF-IDF weighting^55^ before using them for the Random Forest models.

### Nation-of-origin data

Addgene has lab country information for many depositing labs (https://www.addgene.org/browse/pi/). For those missing, publication links were followed to affiliation addresses, and the country of the lab was manually read off the address and cross-checked with a web search. When country information was missing from a lab, if there was conflicting information, and for very rare countries, these classes were dropped and all the corresponding plasmids for that lab were dropped from all three training split sets. No reshuffling of the train, validation and test data occured.

### Lineage network analysis

Lab, country, and ancestry-descendent linkage counts were obtained from the training set and plotted as described above, rank-ordering where specified. Networks were analyzed for size and graph diameter with NetworkX. Lab lineages were obtained from plasmid lineage data by considering the presence of a link from plasmid X from lab A to plasmid Y from Lab B to be a directed edge from Lab A to B. Parallel edges were not allowed and weight was not considered. Self edges were disregarded. The country lineage network was constructed from the lab lineage network, by considering a connection between lab X in country A and lab Y in country B to be a directed edge between country A and B. This time, weights were given as the number of lab-to-lab connections. Parallel edges were not allowed, but self-edges were. For simplicity, the arrows of the directed graph were not shown in the NetworkX visualization. A version of Google PageRank^56^ was computed on the directed, weighted graph with NetworkX.

### Ancestor lab prediction

Due to the earlier biased train-validation-test split which deliberately segregated lineage networks into one of the three sets to minimize ancestry relationships that could lead to overfitting, we reconsidered the dataset for ancestor lab prediction. By definition, an ancestor plasmid and all its descendents will always be in the same set. So, if the ancestor is in the validation set, none of its descendants are available for training.

Therefore, we first parsed the most recent ancestor for each plasmid from the lineage data and assigned each plasmid that ancestor’s lab. We then randomly 80-10-10 resplit the data.

We recognize that there is some potential for meta-overfitting by performing this reshuffle, even though ancestor lab prediction is a unique task from any of the others done so far. However, as this analysis was using a simple model intending to show tractability rather than peak performance, we decided enabling the right train-test-split was worth the chance.

### Interpreting the deteRNNt model

To visualize the hidden states of the model, we first performed inference of the deteRNNt model on the validation set and extracted the activations of the last hidden layer (just prior to the layer which outputs logits). These hidden states were 1000-dimensional. To visualize them, we projected them into two and three dimensions using tSNE^39^ in scikit-learn with default hyperparameters. We colored each point by P(True Lab) as assigned by the model for that example. Note that throughout this section when we refer to model probabilities, we mean the probabilities given by the model after temperature scaling calibration has been applied. The three dimensional interactive plot was made using plotly and labelled with each plasmid’s true lab of origin.

For the analyses of sequence motifs (Fig. 5 b-e) we use the deteRNNt sequence only-model as phenotype information was not available for all plasmids. For the scanning-N ablation analysis (Fig. b,c), we made all possible sequences with a window of 10 Ns inserted and performed standard deteRNNt inference to predict logits. We then generated per-position predicted logits by selecting all the sequences which include a given position as mutated to N, and averaging their logits together. We then apply softmax over each of these position logits to generate per-position predicted probabilities, which are indexed by the relevant labs to visualize.

For the scanning subsequence analysis (Fig. 5 d-e) we made all possible subsequences of given length K and predicted logits for each, as above. Note that these are subsequences in isolation, as if they were sequences from a new plasmid, rather than padded with Ns or similar. For each position in the full sequence, we selected all the subsequences which include that position and averaged together their predicted logits, softmaxing to visualize probabilities as above.

We custom-designed a gene-drive vector using the Akbari germline-Cas9 plasmid AAEL010097-Cas9 (Addgene #100707) as a baseline. We modified it by removing the eGFP sequence attached to Cas9 and replacing it with the dsRed1 sequence within the same plasmid (but removing the Opie2 promoter in the process). We then identified two Cas9 guide RNAs against the *Aedes aegypti AeAct-4* gene^57^ (Genbank Accession Number: AY531223), designed to remove most of the coding region, that are predicted to have high activity using CHOPCHOP (https://chopchop.cbu.uib.no)^58^. These guides were placed downstream of the Akbari identified AeU6a and AeU6c promoters (Addgene #117221 and #117223)^59^. We also included somArchon-GFP from pAAV-Syn-SomArchon (Addgene #126941, deposited by Ed Boyden’s group) as a non-Akbari derived sequence. The Cas9-dsRed1_guide cassette_somArchon-GFP payload was flanked by 500bp homology arms (upstream of 5’ guide and downstream of 3’ guide).

To analyze the results of our ablation and subsequence analyses we indexed out the positions in the sequence with the most extreme changes in predicted probability and manually examined these regions in Benchling. We performed automated annotation, used BLAST^60^, and searched various repositories for the highest-ranked fragments in order to identify restriction sites, primers sites and other features.

### Baselines

The comparison with BLAST was performed using the blastn command line tool from NCBI^60^. At a high level, we can consider the BLAST baseline to be a nearest-neighbor algorithm, where the blast e-value is used to define neighbors in the training set. For each of the lab-of-origin and nation-of-origin prediction tasks, a fasta file of plasmid sequences from the training set was formatted as a BLAST database. Then, each test set was blasted against this database with an e-value threshold of 10. The resulting training set hits were sorted by e-value, from lowest to highest, and used to look up the training set labels for each sequence. For top 1 accuracy, the lowest e-value sequence class was compared to be the true class. For top 10 accuracy, an example was marked correct if one of the labels of the lowest 10 e-value hits, after dropping duplicate hits to the same lab, corresponded to the correct test set label. Dropping duplicates ensured that the BLAST baseline was permitted up to 10 unique lab “guesses”, which is necessary because occasionally the top-k ranked sequence results all have the same lab label. To perform the nation-of-origin and U.S. vs foreign comparison, the same blast results were filtered to drop the U.S. or binarized so the U.S. was the positive class, respectively.

The comparison with the Convolutional Neural Network (CNN) method copied the architecture and hyperparameters reported in Nielsen & Voigt (2018)^23^. The model was implemented in PyTorch. We trained for 100 epochs, as reported in previous work^23^, on an Nvidia K80 GPU using the Amazon Web Services cloud p2.xlarge instance. Training converged on a validation score of 57.1% and appeared stable (Supplementary Fig. 6). After training for this duration, we saved the model and evaluated performance on the held-out validation and test sets. Test-set performance was 50.2%, which was very near the previously reported accuracy of 48%^23^, leading us to conclude that this model’s performance was reproducible and robust to increases in both the number of labs and number of plasmids in our dataset compared to Nielsen & Voigt (2018) (827 vs. 1314 labs, 36,764 vs. 63,017 plasmids)^23^. In other words, this replication of the architecture, hyperparameters and training procedure of previous work with everything held constant to the best of our knowledge except the dataset, suggests that the effect of having more examples (typically associated with an easier task) and more labs to distinguish between (typically associated with a more difficult task) approximately cancel out, or perhaps net to a very weak (2%) difference in the difficulty of our dataset.

In lab-of-origin, U.S. vs foreign, nation-of-origin, and ancestor lab prediction, we show comparisons with a baseline based on guessing the most abundant class, or classes (in the case of top 10 accuracy) from the training set. We also show the frequency of success based on uniformly guessing between the available labels (1/ number of categories).

## Supplementary Information

**Supplementary Table 1.**
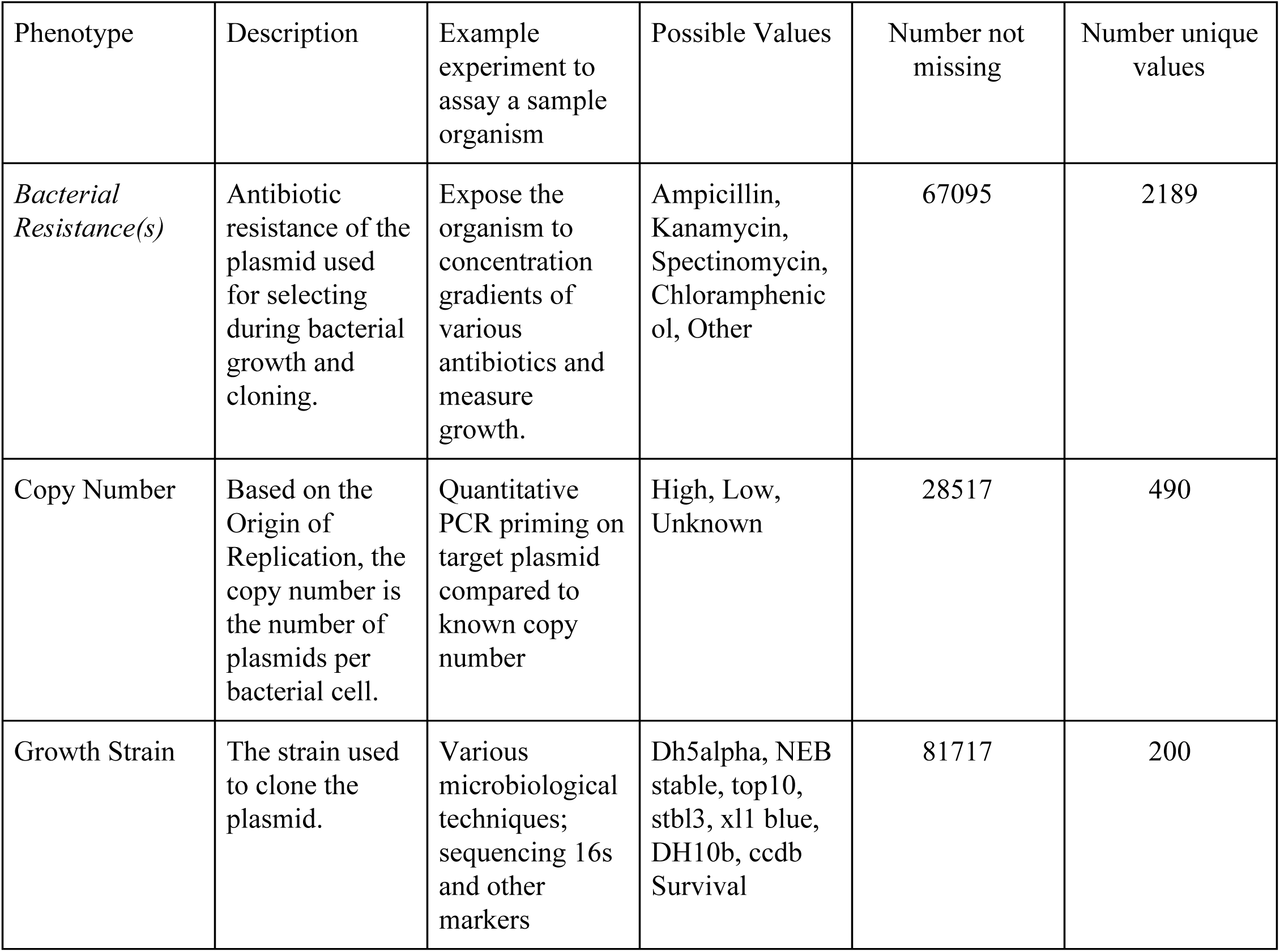
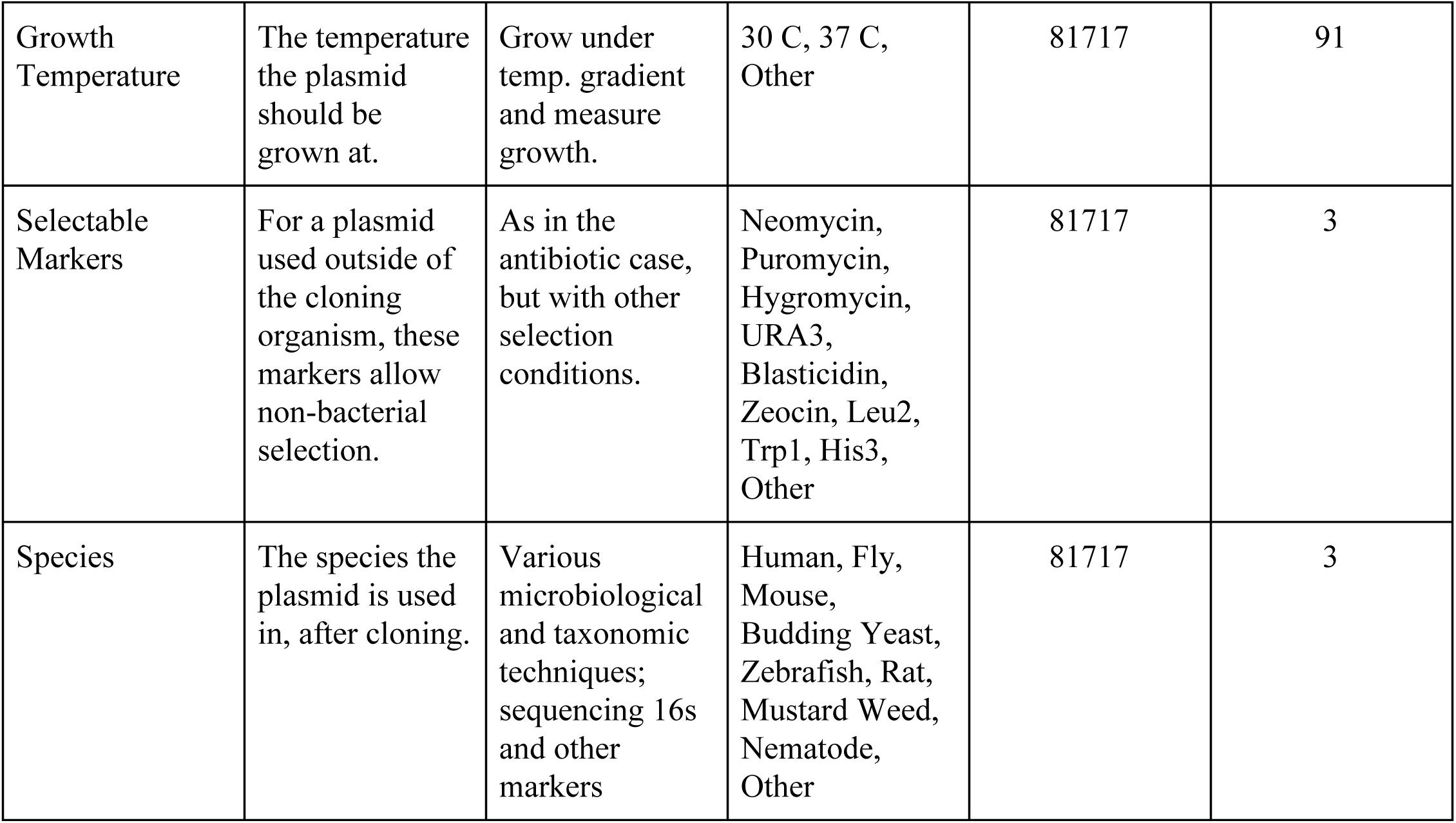
Simple phenotypic information inferred from Addgene. Note that in the envisioned deployment scenario some of these characteristics may be unavailable or uniformative, for example the growth strain used for cloning is not available if you have a sample of the chassis orgamism instead of cloning strain. On the other hand, many additional and more informative phenotypic characterizations are possible, and could be incorporated into the model if a sufficiently large and representative dataset of measurements and lab-of-origin are assembled.

**Supplementary Figure 1.**
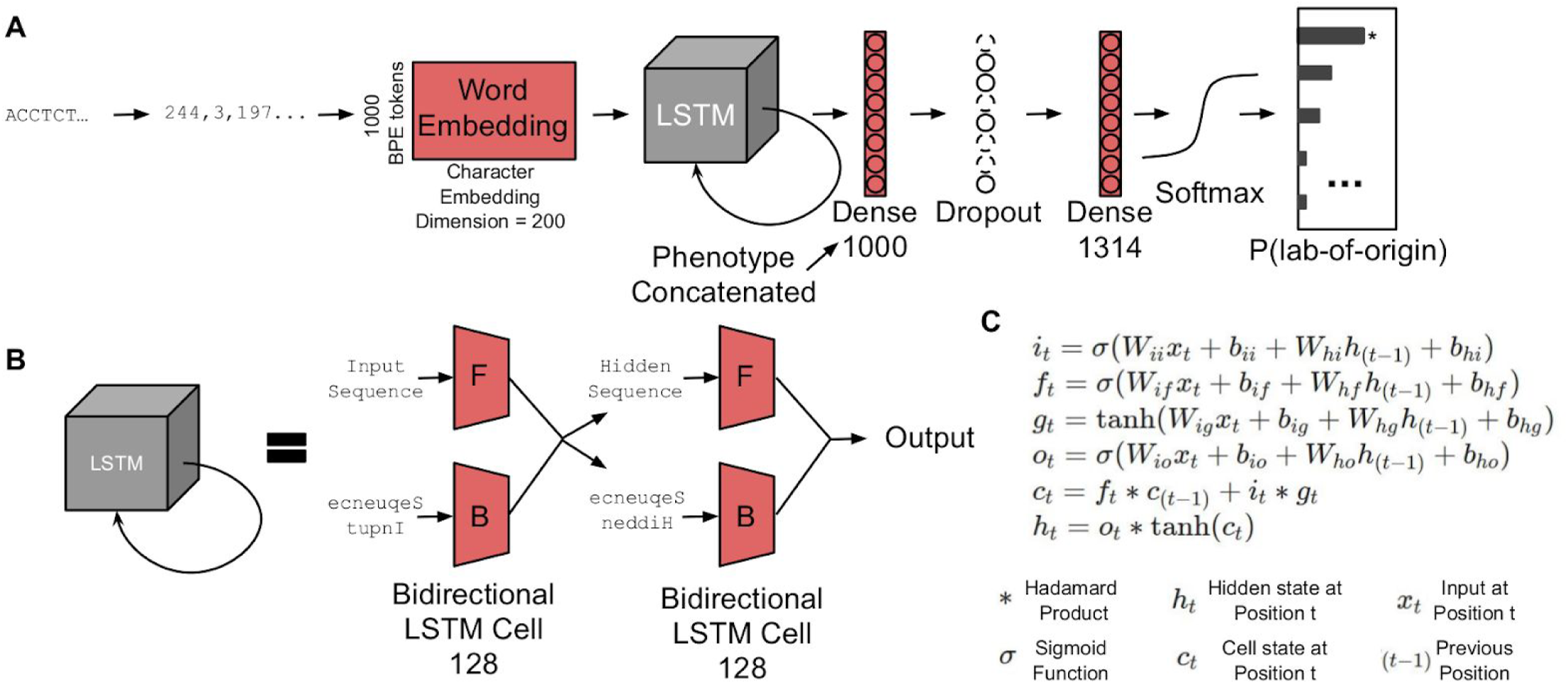
Model architecture. (**A**) From left to right: a DNA sequence is Byte-Pair-Encoded to integers, embedded into a vector space, passed through a 2-layer bi-directional LSTM, to a 1000-d fully connected layer with dropout = .5, another fully connected layer which outputs 1314 dimensional logits, one for each class, which is softmaxed to produce a prediction vector summing to 1. (**B**) The LSTM. Input sequence is processed forward (F) and backward (B) by LSTM cells. The previous layer’s hidden states are concatenated, and as before both the forward and backward sequence are processed. The output is the concatenation of the final hidden state from forward and backward cells. (**C**) Mathematical definition of an LSTM cell. At each position along a sequence, the output of the cell is defined by the value of the input sequence (x_t_) and a recurrent relationship with the previous step, captured in a hidden state and cell state (*h*_*(t-1)*_ *c*_*(t-1)*_). Typically, *i*_*t*_, *f*_*t*_, *g*_*t*_, *o*_t_, are called the input, forget, cell and output gates, respectively. For motivation and a full mathematical treatment, please see Hochreiter and Schmidhuber (1997)^61^.

**Supplementary Figure 2.**
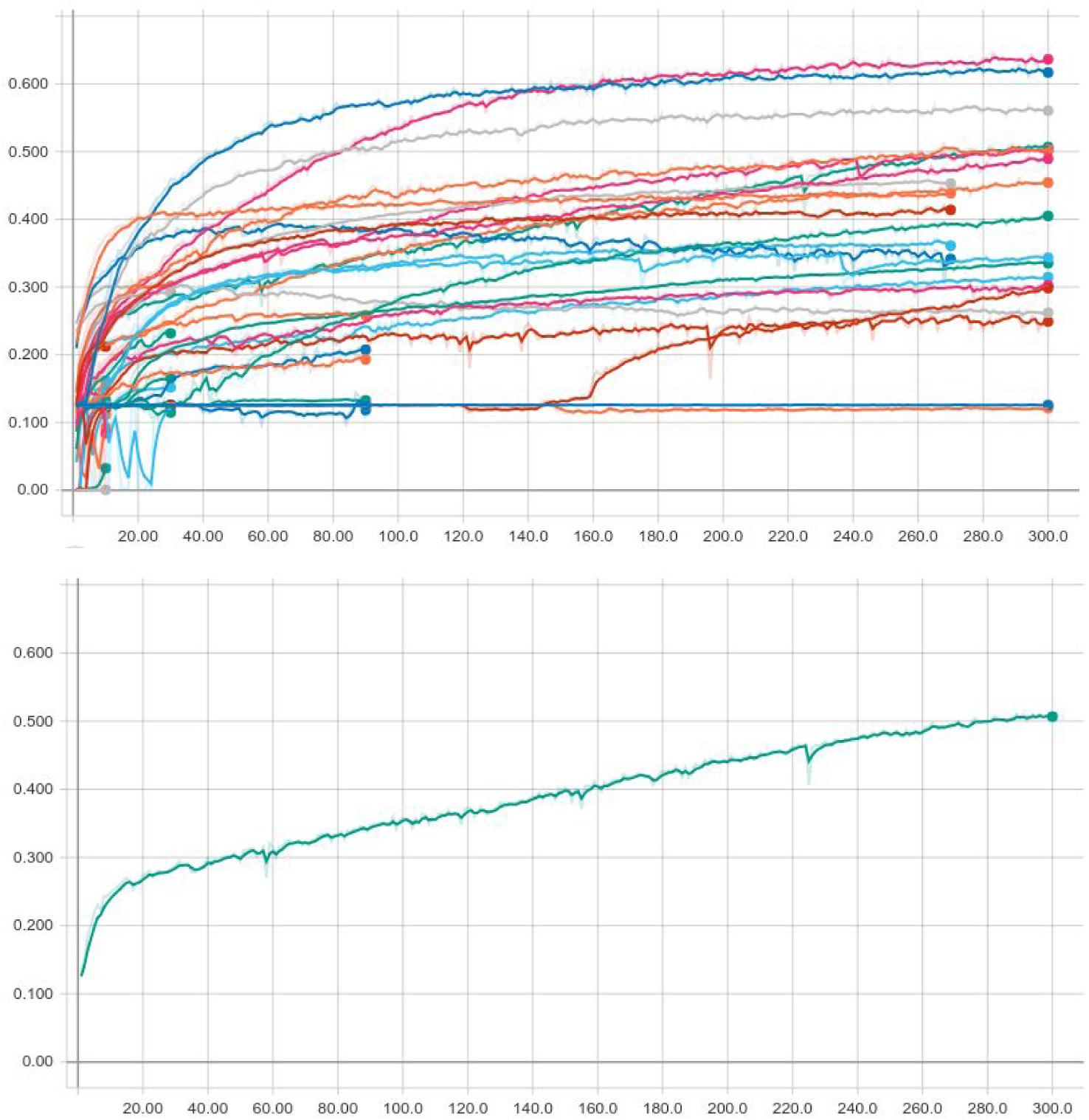
Accuracy curves of Asynchronous Hyperband hyperparameter search. Validation accuracy on the Y axis, and number of Hyperband steps on the X axis. The entire collection of variants (top) is compared to the selected model (bottom).

**Supplementary Figure 3.**
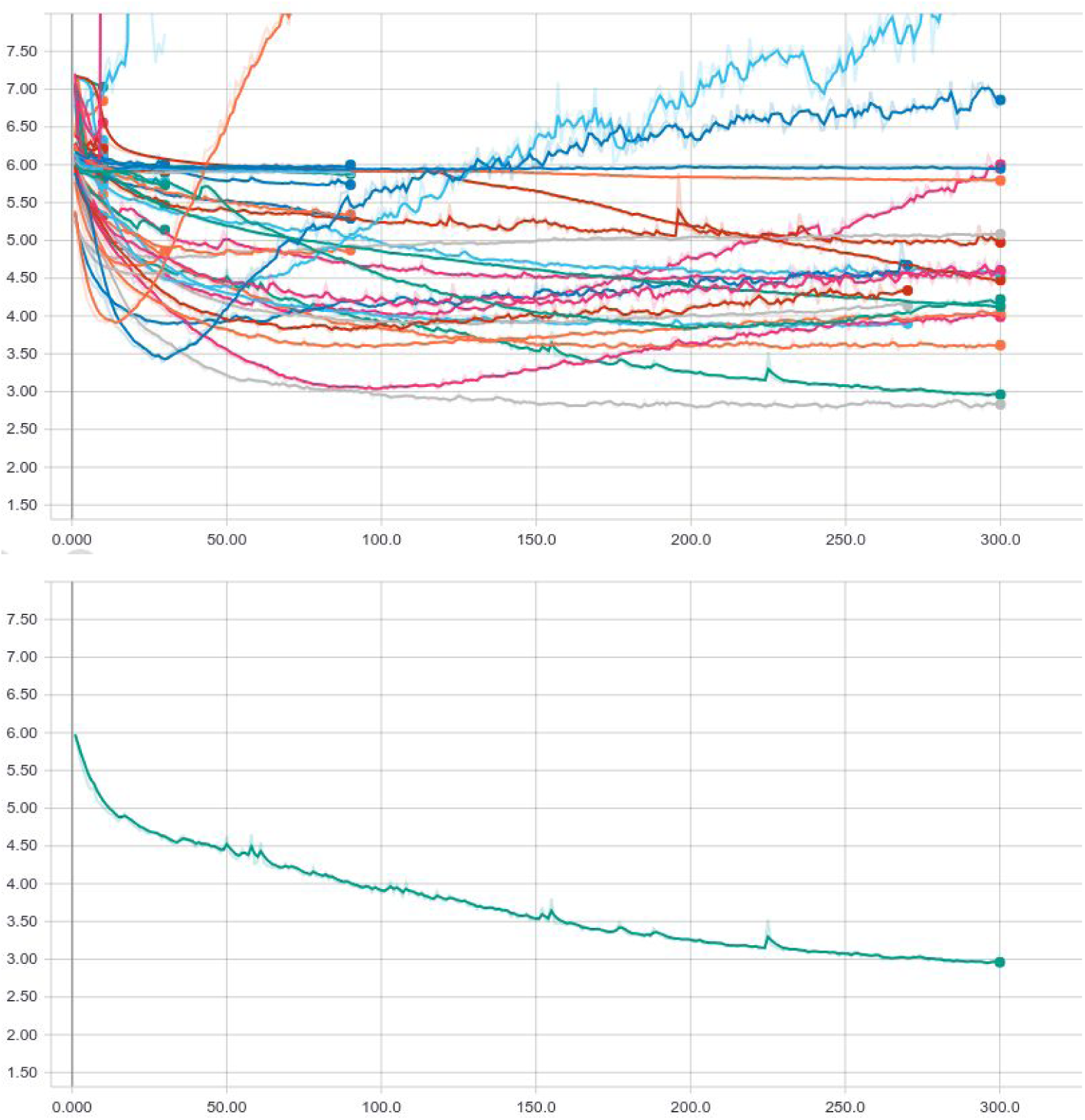
Loss curves of Asynchronous Hyperband hyperparameter search. Validation set cross entropy loss on the Y axis, and number of Hyperband steps on the X axis. The entire collection of variants (top) is compared to the selected model (bottom).

**Supplementary Table 2.**
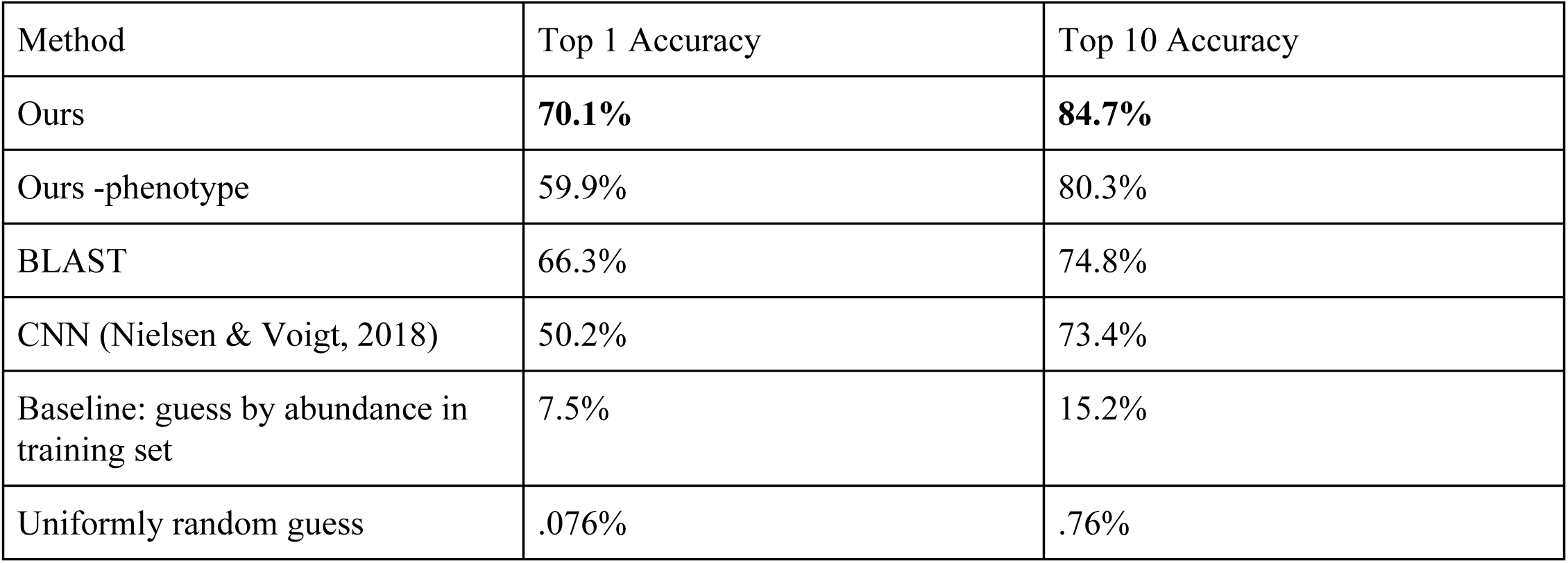
Lab-of-origin attribution accuracy on the held-out test set.

**Supplementary Table 3.** Nation-of-origin attribution accuracy on the held-out test set.

| Method | Top 1 Accuracy | Top 10 Accuracy |
| --- | --- | --- |
| Random Forest | **75.8%** | **96.7%** |
| BLAST | 70.3% | 86.7% |
| Baseline: guess by abundance in training set | 16.1% | 83.8% |
| Uniformly random guess | 3.0% | 30.3% |

**Supplementary Table 4.** Ancestor lab attribution accuracy on the held-out test set.

| Method | Top 1 Accuracy | Top 10 Accuracy |
| --- | --- | --- |
| Random Forest | **87.0%** | **96.5%** |
| Baseline: guess by abundance in training set | 13.6% | 45.0% |
| Uniformly random guess | .5% | 5.3% |

**Supplementary Figure 4.**
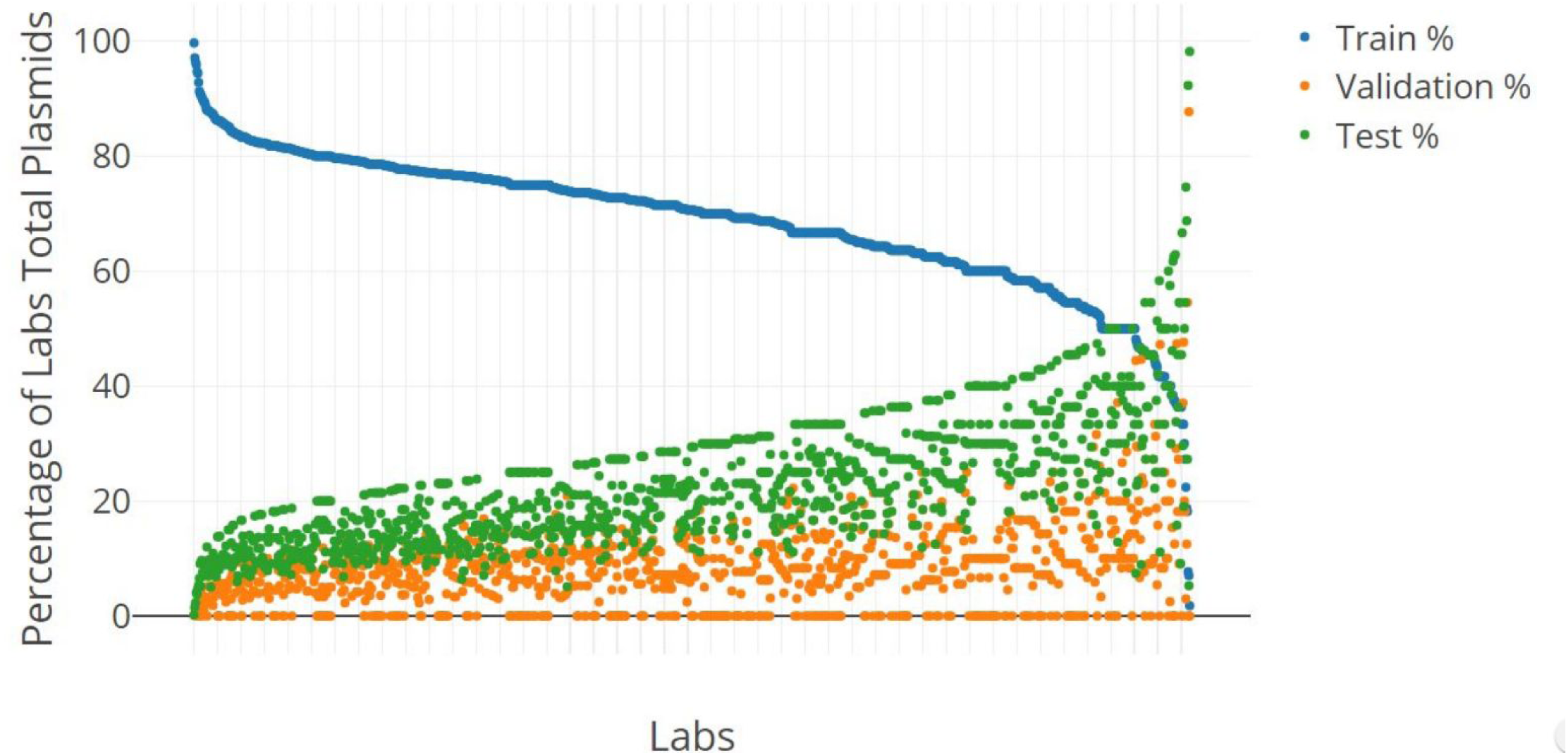
Lab distribution after train-test-validation split. Each vertical sums to 100%. The validation set points (orange) hit 0% abundance because there was no rule that the validation set must have a certain number of plasmids per lab.

**Supplementary Figure 5.**
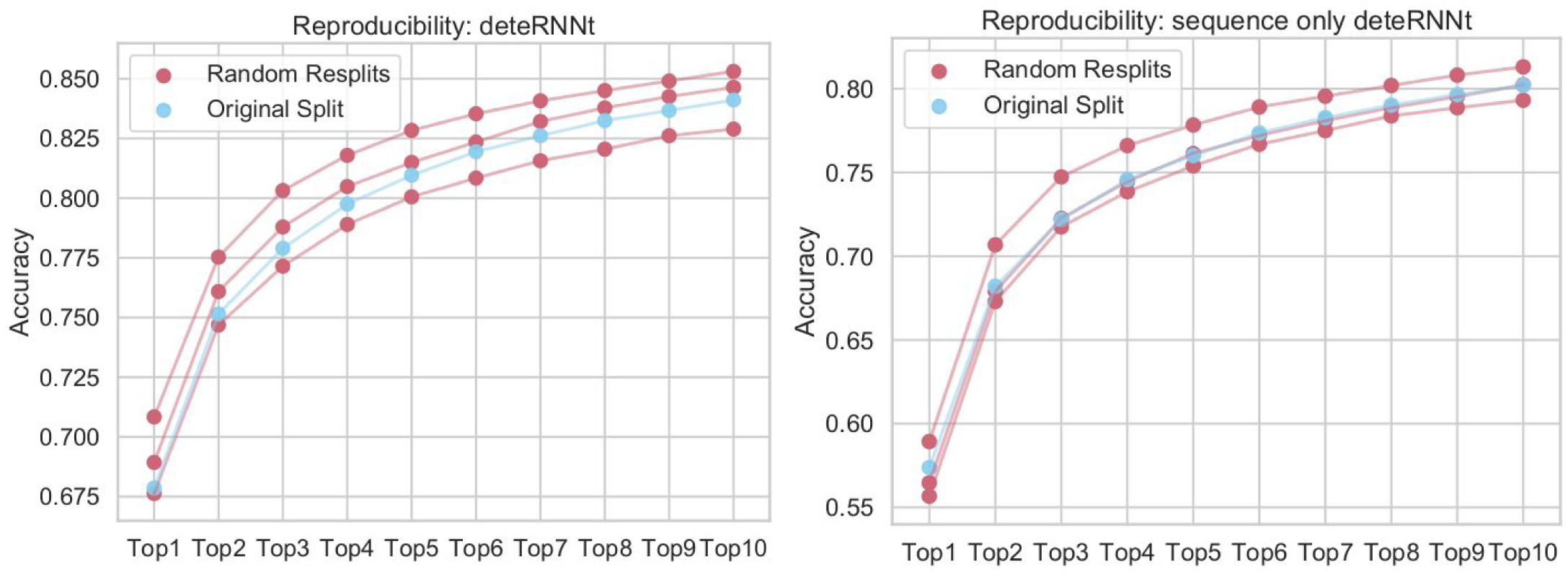
Reproducibility of random data splits. Three full resplits of the data show on-par performance with the original split, using the same hyperparameters but different random seeds and input data. X axis shows TopN accuracy for N in [1-10]. Y axis shows test set accuracy for the splits corresponding test set. The original split (blue) has a different random seed and slightly different training time from the model presented in the main text but the same data.

**Supplementary Figure 6.**
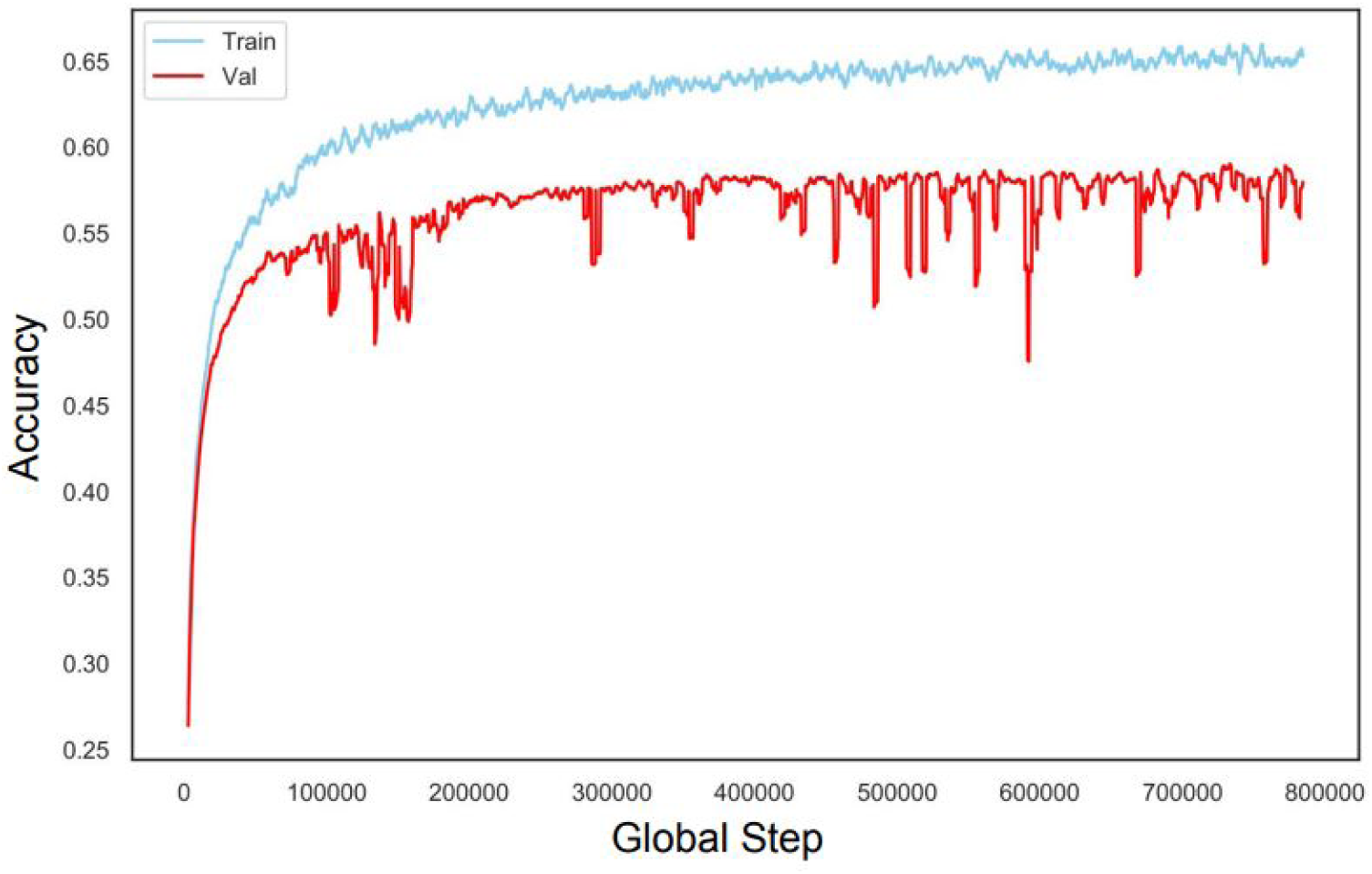
Training and validation curves of the CNN model on Addgene lab-of-origin data. Training continues to 100 epochs, or 787,800 steps on our data.

